# Seroprevalence of anti-microbial antibodies in the normal healthy population with implications in chronic diseases

**DOI:** 10.1101/693655

**Authors:** Peilin Zhang, Lawrence M. Minardi, J. Todd Kuenstner, Steve M. Zekan, Rusty Kruzelock

## Abstract

We have previously discovered a panel of anti-microbial antibodies from patients with Crohn’s disease (CD) and Sjogren’s syndrome (Sjo). We have also demonstrated the increase of these anti-microbial antibodies in other autoimmune disease such as rheumatoid arthritis, systemic lupus erythematosus and multiple sclerosis in a small number of cases. The seroprevalence of these antibodies in the normal healthy population is unknown. We set to survey the normal population for these anti-microbial antibodies. We collected 288 blood samples from the donor units of the leukocyte-reduced red blood cells from the American Red Cross, and examined the presence of the anti-microbial antibodies in these blood samples using direct ELISA assays established in our laboratory using the recombinant microbial protein antigens. Our results showed that the prevalence of RPOB antibody in the normal blood donor population is 2.4% (7 positive of 288 samples). The prevalence of EF-G antibody is 4.2% (12 positive of 288 samples), ATP5a 5.2% (15 positive), Hsp65 2.8% (8 positive), EF-Tu 5.6% (16 positive), and NMPC 4.2% (12 positive). Meanwhile, in 109 patients with Crohn’s disease and 28 patients with Sjogren’s syndrome, these anti-bacterial antibodies are significantly increased (p<0.001). These results indicate that the specific anti-microbial antibodies within the normal general population are uncommon, but frequent in chronic disease states. The presence of increased anti-microbial antibodies in the blood of patients but not in normal controls can serve as biomarkers for chronic diseases such as Crohn’s disease and Sjogren’s, and their presence indicates abnormal B-cell/plasma cell function in response to the commensal/pathogenic microbes. Since the antigens were derived from the common microbes present on the surface of the normal population, the antimicrobial antibodies in patients with diseases but not in the normal population suggest a deficient clearance of the microbes from the circulation by the innate immunity system in chronic diseases. These results also raise questions of bacterial vaccination using whole bacterial extracts as these anti-bacterial antibodies appear pathogenic rather than protective, offering fresh thinking in designing bacterial vaccines as preventive or therapeutic measures in chronic diseases such as Crohn’s disease and Sjogren’s Disease.

## Introduction

Specific anti-microbial antibodies in the circulation have been previously described and successfully used for the diagnosis of Crohn’s disease (CD), ulcerative colitis and irritable bowel disease using the inflammatory bowel disease (IBD) panel from Prometheus Laboratory (San Diego, CA)^[1–6]^. Our previous studies showed a serologic antibody panel against a group of novel microbial proteins and this panel of specific antibodies against the microbial proteins is significantly elevated in patients with Crohn’s disease (CD) and Sjogren’s syndrome (Sjo) in comparison to normal healthy controls using indirect ELISA assays ^[7 8]^. Three of the six previously identified anti-microbial antibody markers, DNA directed RNA polymerase subunit B (RPOB), Elongation factor G (EF-G) and ATP synthase subunit alpha (ATP5a), are derived from *Staphylococcus aureus (S. aureus & Pseudintermedius)*. Heat shock protein 65 (Hsp65) is derived from *Mycobacterium tuberculosis (MTB)* or other non-tuberculous mycobacteria *(Mycobacterium avium paratuberculosis (MAP) and Hominissuis, (MAH)*), and elongation factor Tu (EF-Tu), and outer membrane porin protein (NMPC) are from *Escherichia coli (E. coli)*^[8 9]^. These microbes are normally present on the surface or in the gut of the human host or in the environment in the case of non-tuberculous mycobacteria, and are considered commensal bacteria or non-pathogenic bacteria/mycobacteria under normal circumstances. Since the specific anti-microbial antibodies are directed to the commensal bacteria in the human body, the presence of these anti-microbial antibodies in the patients raises questions about the patient’s own genetic susceptibility, rather than implicating the virulence of the microbes.

Alternatively, the microbes change their virulence in the patient population and these microbes are different to certain degrees from those colonizing the normal human population. To address this question, the seroprevalence of these anti-microbial antibodies in the normal healthy population is important. In the setting of physicians’ offices, it is difficult to obtain the “normal” control population since the “normal” asymptomatic human being will not visit the physicians’ offices, thus we turn to the blood donor population from the American Red Cross.

Advancement of microbiome study of the normal healthy human being and the patients with Crohn’s disease (CD) indicates the presence of a large number of aerobic and anaerobic microbes including fungi within the human gut, and this large quantity of microbes maintains a systemic balance for the human health and metabolism^[10–13]^. Imbalance of these microbes occurs in a variety of chronic disease conditions (dysbiosis)^[13]^. How these gut microbes interact with the host immune system is gradually emerging ^[14–16]^. However, the anti-microbial antibodies in many chronic diseases have not been extensively studied, and how and why the anti-microbial antibodies are generated in the patients with various disease conditions is poorly understood.

We have established the direct ELISA assays for anti-microbial antibodies using the recombinant microbial protein antigens. Direct ELISA assay can simplify the testing process significantly, and increase the sensitivity, specificity and reproducibility. We obtained 288 normal donor blood units from the local Red Cross office and surveyed the prevalence of the anti-microbial antibodies in these normal blood donors to assess the presence and the distribution of these anti-microbial antibodies. We believe this effort is an essential step towards understanding the mechanism of the pathogenesis of many chronic diseases in relation to the microbiome within the human body.

## Material and methods

### 1, Direct ELISA assays

#### A, Recombinant protein biosynthesis

Recombinant microbial protein synthesis was performed by Genscript Corporation (http://www.genscript.com/) through contract work commercially. The microbial proteins and the corresponding Genbank numbers were identified at PZM Diagnostics through a series of identification and mass spectrometry as described previously ^[8 9]^. The genes encoding for the microbial proteins were synthesized chemically, and cloned into pUC57 bacterial expression vectors with His-tag using proprietary technology at Genscript Corp. The microbial proteins were expressed in the *Escherichia coli* expression system, and purified through His-tag columns. The final microbial proteins were analyzed by Western blot analysis using anti-His-tag antibody and Coommassie blue stain of SDS-polyacrylamide gel for quality assurance. The synthesized recombinant proteins are delivered on dry ice to us and the protein concentration is adjusted at 10 μM with phosphate buffered saline (PBS) with 20% glycerol for storage.

#### B, Establishment of direct ELISA assay

Our direct ELISA assays for human antibody testing is based on the Biolegend protocol with modification (https://www.biolegend.com/protocols/sandwich-elisa-protocol/4268/). Direct ELISA assays for human antibody levels against various recombinant microbial antigens were established based on a series of antigen titration, and stringent detergent levels. The optimal concentration of the recombinant antigens for coating the 96-well plates was determined to be in the range of 0.1 to 1.0 nmol (1-10 ng/ml), and the plate coating was optimal at 4°C over night. The carbonate coating buffer, blocking agents and concentration, washing buffer, substrate (TMB) and stop buffer were all based on the protocol from BioLegend (San Diego, CA). The data was collected by using the VersaMax ELISA plate analyzer from Molecular Device at OD450 TMB using the endpoint protocol. The data was analyzed and plotted by using various components of R-statistics (Rstudio) package (http://statistics4everyone.blogspot.com/).

### 2, Blood donor units and sample processing

Red Blood Cells (RBCs) from the American Red Cross consist of erythrocytes concentrated from whole blood donation or collected by apheresis ^[17]^. Based on the guideline from the American Red Cross, the RBC units contain citrate anticoagulant and usually one of several types of preservative solutions. Depending on the preservative-anticoagulant, the hematocrit (Hct) of RBCs is about 55-65% for additive solutions (AS), AS-1, AS-3, AS-5, AS-7 and about 65-80% for citrate-phosphate-dextrose adenine solutions, CPDA-1, CPD, CP2D. RBCs contain 20-100 mL of donor plasma, usually less than 50 ml, in addition to preservative and anticoagulant. The typical volume of AS RBCs including additive solution is 300-400 mL. Each unit contains approximately 50-80 grams of hemoglobin (Hgb) or 160-275 mL of red cells, depending on the Hgb level of the donor, the whole blood collection volume, and the collection and processing methods ^[17]^. Leukocyte-reduced RBCs must retain at least 85% of the original RBCs. Each unit of RBCs contains approximately 250 mg of iron, almost entirely in the form of Hgb. This varies depending on the original volume and concentration of the unit. There are a few tubal segments attached to the units containing the same RBCs, and these tubal segments are to be used for quality assurance testing or additional studies once the blood unit is transfused. The segments of the donor blood units were obtained from the local Red Cross offices after the blood units were transfused to the patients. One segment from each unit of blood was obtained and processed as the following. The blood storage buffer from the collection and the processing is typically proprietary additive solutions AS1, AS3 or AS7 containing phosphate-based buffer with added adenosine and other components in the US. These storage buffers are compatible with most typical antibody testing methods. Based on the general calculation, the plasma components containing the antibodies in the blood unit is estimated to be less than 10% (1:10 dilution). One segment of the blood unit contains 0.5 cc leukocyte-reduced red blood cells in AS buffer. Assuming the red cell concentration in the storage buffer is 50% (250 microliters), and the plasma component in storage buffer is 50% (250 microliters). After the segment was obtained, one end of the segment was cut with a scissors and the open end was emptied in a 1.5 ml microfuge tube. Then the other end was cut with a scissors and the blood in the tube was released to the tube entirely. An aliquot of 0.5 ml normal saline was used to wash the remaining blood from the open segment, resulting in approximately 1.0 ml red cells with the buffer (the final dilution is estimated to be 1:40 of the original plasma). The microfuge tube with blood was centrifuged at 10,000 g for 1 minute, and the supernatant from the tube (50 ul) was directly used for direct ELISA assays.

The plasma samples from the patients with Crohn’s (CD) and Sjogren’s (Sjo) syndromes were previously collected through commercial testing services at PZM Diagnostics, and the clinical characteristics of these patients were previously described ^[9]^. The plasma samples were re-tested using the newly established direct ELISA methods using recombinant microbial protein antigens. The plasma samples were diluted 1:50 using normal saline with 5% BSA and the diluted plasma samples were used for direct ELISA assays. The control group was from the physicians’ offices that we have collected over the years with no evidence or history of Crohn’s disease (CD) and Sjogren’s syndrome (SJO).

## Results

### 1, Seroprevalence of anti-bacterial antibodies in normal healthy individuals from the donor population of the Red Cross

A total of 288 donor samples were used for standard direct ELISA assay for detection of the antibodies within the donor samples against the recombinant RPOB, EF-G, ATP5a, HSP65, EF-Tu and NMPC. The blood donors of the Red Cross have been screened under strict regulations in the US and the segments of the donor units are the identical blood units that have been transfused to the patients. The segments are usually retained at the hospital blood banks for 3 months at 4° C before being discarded after transfusion. The segment samples were tested using the newly established direct ELISA assays for antibodies against RPOB, EF-G, ATP5a, Hsp65, EF-Tu, and NMPC as described in the above sections. The data of the entire results is summarized in Table 1 including the mean, standard deviation (SD), median, high and low values for each analyte, 95% confidence interval for the mean, cutoff value for each analyte, and the total number and percentage of positive samples in this population based on the cutoff values. The mean for RPOB antibodies in 288 Red Cross donors is 0.184 with standard deviation (SD) 0.116. The cutoff value for RPOB was 0.42 (Mean + 2 x SD). There were a total of 7 positive samples for RPOB based on the cutoff value of 0.42, representing 2.4% within this Red Cross donor population. Similarly, the cutoff value for EF-G was 0.48, and there were 12 positive samples, representing 4.2%. There were 15 positive samples for ATP5a (5.2%), 8 positive samples for Hsp65 (2.8%), 16 positive for EF-Tu (5.6%) and 12 positive for NMPC (4.2%).

**Table 1:** Performance of six anti-microbial antibodies in Red Cross donor population

| **<u>N=288</u>** | **<u>RPOB</u>** | **<u>EF-G</u>** | **<u>ATP5a</u>** | **<u>Hsp65</u>** | **<u>EF-Tu</u>** | **<u>NMPC</u>** |
| --- | --- | --- | --- | --- | --- | --- |
| Mean | 0.184 | 0.21 | 0.21 | 0.113 | 0.208 | 0.186 |
| Median | 0.17 | 0.174 | 0.18 | 0.12 | 0.164 | 0.163 |
| SD | 0.116 | 0.136 | 0.128 | 0.107 | 0.129 | 0.106 |
| 95% confidence interval for mean | 0.170 -- 0.200 | 0.195 -- 0.224 | 0.196 – 0.223 | 0.100 -- 0.130 | 0.194 -- 0.221 | 0.173 – 0.200 |
| High | 0.885 | 0.98 | 0.783 | 0.85 | 0.726 | 0.84 |
| Low | 0.04 | 0.04 | 0.04 | 0.036 | 0.04 | 0.04 |
| Cutoff (mean + 2 x SD) | 0.42 | 0.48 | 0.47 | 0.33 | 0.47 | 0.4 |
| Total positive (percentage) | 7 (2.4%) | 12 (4.2%) | 15 (5.2%) | 8 (2.8%) | 16 ( 5.6%) | 12 (4.2%) |

### 2. Anti-microbial antibodies in Crohn’s disease and Sjogens syndrome

We have collected 109 plasma samples from Crohn’s patients (CD) and 28 from Sjogren’s syndrome (Sjo). A portion of these samples were tested for the same antimicrobial antibodies using indirect ELISA assays with specific capturing antibodies and whole bacterial extracts as previously described^[8 9]^. Currently we re-tested these samples using the newly established direct ELISA assays and specific recombinant microbial protein antigens. We have collected 156 controls from the clinics and physicians’ offices, and these controls from the clinics and physicians’ offices were used as controls for Crohn’s disease but these controls were not entirely normal healthy individuals as a variety of chronic conditions such as rheumatoid arthritis, thyroid diseases, chronic fatigue syndrome, multiple sclerosis, etc are known for these controls. These controls are different from those obtained from the Red Cross blood donors, and as we have noted these chronic conditions are invariably related to abnormal interaction between the human host and its colonizing microbes. As shown in Figure 1, there is a significant increase of all the anti-microbial antibodies in Crohn’s disease (CD) and Sjogren’s syndrome (Sjo) in comparison to the Red Cross blood donors or other controls from the physicians’ offices. (p<0.001 for all six antibodies). The sensitivity of the panel test for Crohn’s (CD) patients ranges from 80.3% to 95.1% (RPOB 90.2%, EF-G, 95.1%, ATP5a 95.0%, Hsp65 90.0%, EF-Tu 80.3%, NMPC 91.8%). There is also a significant increase of all antibodies in the control population from the physicians’ offices in comparison to the Red Cross donor population (p<0.01 for all six antibodies). In a comparison of Crohn’s disease (CD) and Sjogren’s syndrome (Sjo), the results were mixed for all the six specific anti-microbial antibodies (Figure 2). The levels of anti-microbial antibodies for RPOB, EF-Tu and NMPC from the blood of Sjogren’s (Sjo) patients were significantly higher than those of Crohn’s disease (CD) (p<0.05, unpaired T test) but the remaining three were not significantly different (p>0.05).

**Figure 1:**
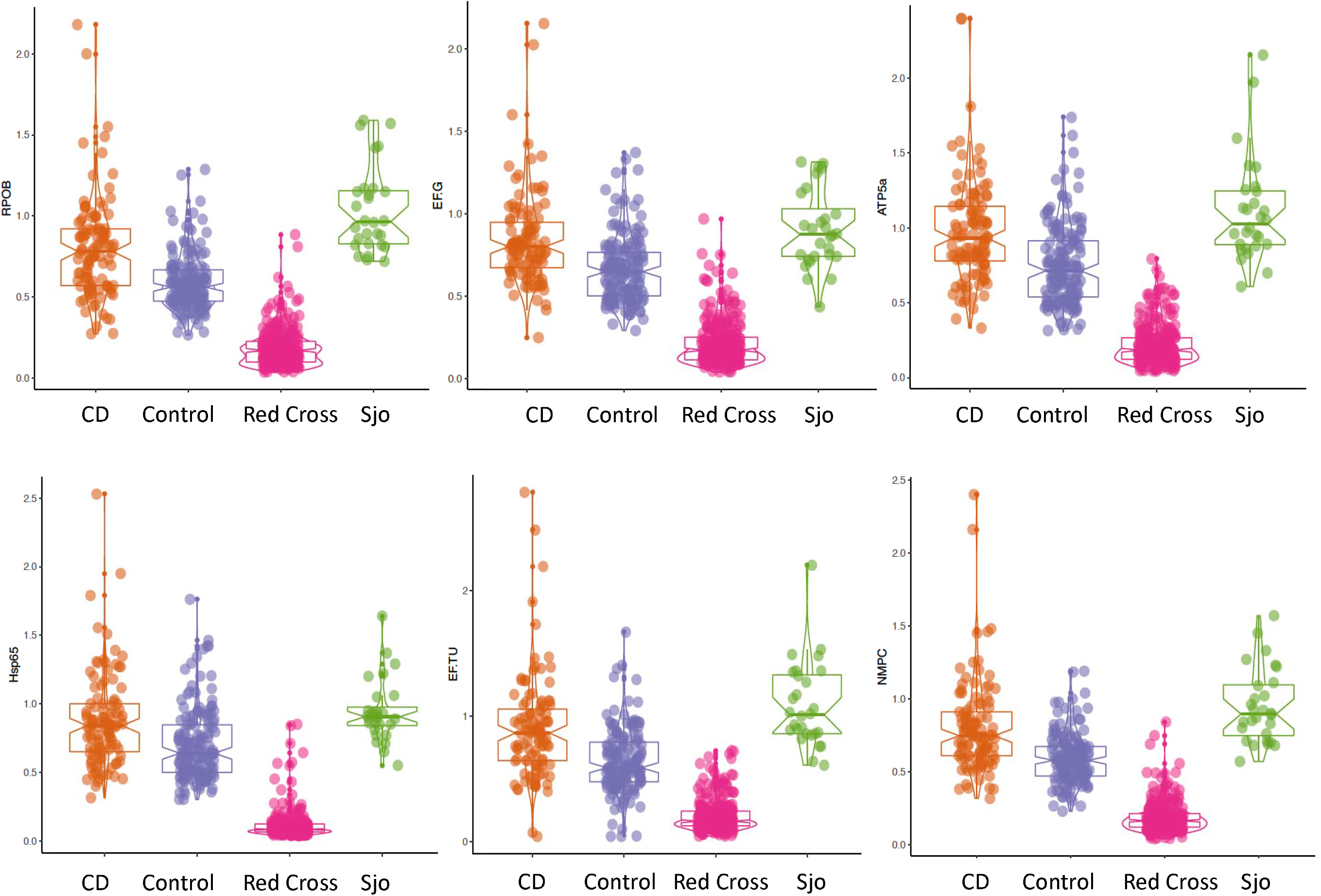
Six anti-microbial antibodies in normal and disease populations. The Y-axis represents the raw OD450 reading using the direct ELISA assays against 6 recombinant microbial antigens. N=109 (CD), N=28 (Sjogren’s syndrome), N=156 (Controls for CD from physicians’ offices), N=288 (Red Cross donor population).

**Figure 2.**
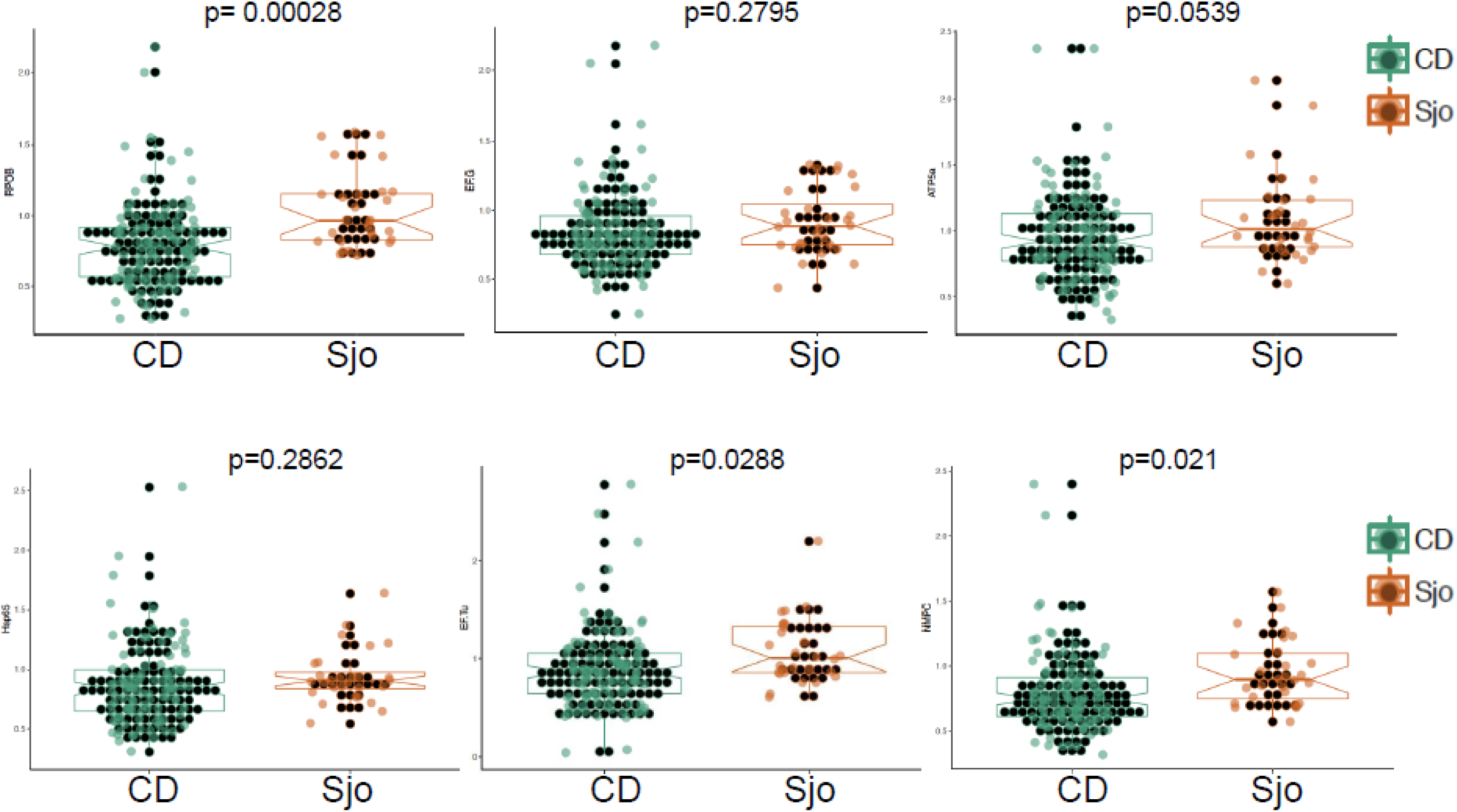
Comparison of anti-microbial antibodies against 6 recombinant microbial antigens using direct ELISA assays in Crohn’s disease and Sjogren’s syndrome. N=109 (CD), N=28 (Sjo, Sjogren’s syndrome).

### 3. Receiver operating characteristic (ROC) curve

ROC curve analysis was performed for all the samples including the controls and the Red cross donors as negative for diseases, and all the samples from Crohn’s (CD) and Sjogren’s (Sjo) patients as positive for diseases using the ROC curve analysis of the R-package ^[18]^ (Figure 3) (http:// http://statistics4everyone.blogspot.com/). The area under the curve (AUC) for RPOB was determined to be 0.899, AUC for EF-G was 0.917, AUC for ATP5a 0.901, Hsp65 0.887, EF-Tu 0.915, and NMPC 0.906.

**Figure 3.**
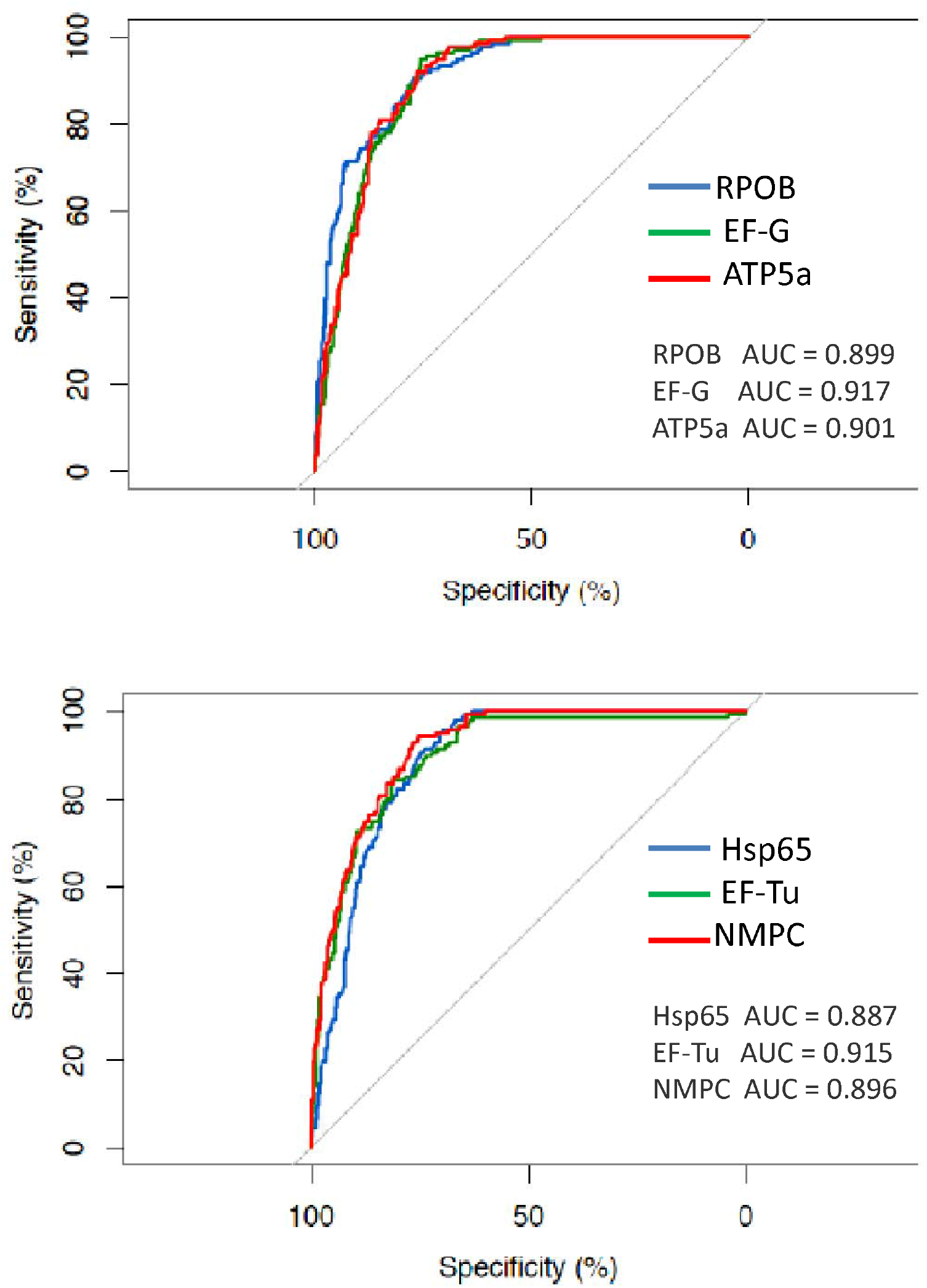
Receiver operating characteristics (ROC) curve demonstrating each of the anti-microbial antibody in Crohn’s disease and Sjogren’s syndrome in comparison to the normal controls including the Red Cross donor population and the controls from the physicians’ offices.

**Figure 4.**
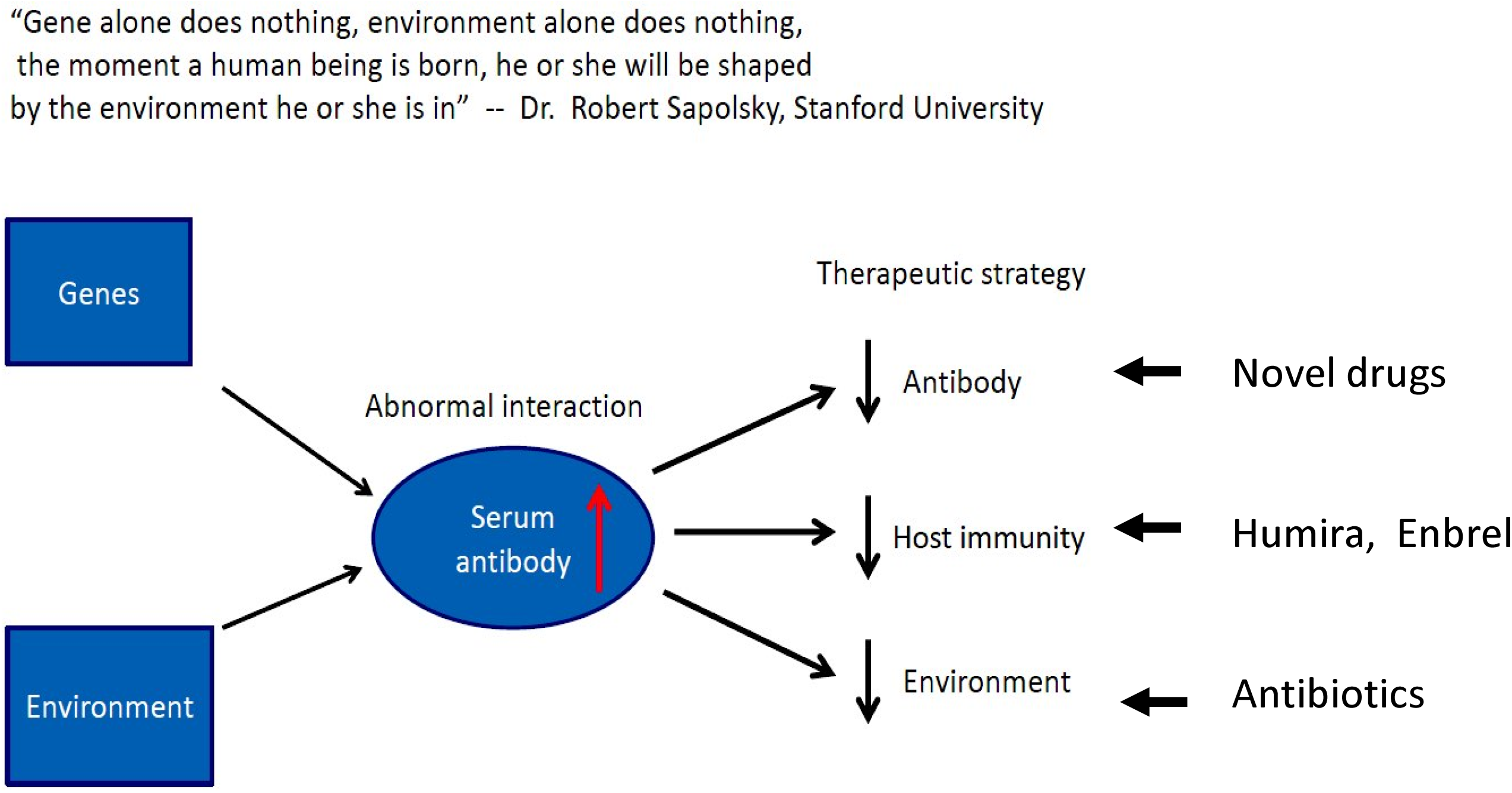
Schematic representation of interaction between the human host and its colonizing microbes, and potential strategy to counter the abnormal interactions in chronic diseases.

## Discussion

Anti-microbial antibodies have been used for diagnosis of microbial infection since George-Fernand Widal over a century ago (Widal test) ^[19]^. The basic principle and the methodology remains similar over the last century, and many clinical diagnostic tests for viral and bacterial infections are still in practical use today. Essentially, the invading microbes are considered “foreign” and these microbes elicit the human immune response to produce neutralizing/opsonizing antibodies to stop the pathogenic invasion. Activation of T-cell/B-cell/plasma cells in response to invading pathogens is considered “adaptive immunity” in contrast to the “innate immunity” consisting of the physical barriers, phagocytic systems and complement systems.

We have surveyed the normal population of blood donors from the Red Cross for the presence of anti-microbial antibodies to determine the seroprevalence of these antimicrobial antibodies. Blood donors of the Red Cross are normal healthy individuals that have been screened for infectious diseases, and this normal population is different from other control patients we have collected for other studies. Our results showed that under the normal conditions in the general population the anti-microbial antibodies are uncommon (2 - 5.6% as shown in Table 1), and this low prevalence of anti-microbial antibodies in the normal population is consistent with the classic concept of human immunity that the bacterial clearance is through the innate immunity, and the adaptive immunity (B- or T-cell activation) is only required to compensate the deficiency of the innate immunity in the clearance of bacterial invasion ^[20]^. There is also a large quantity of immunoglobulins in circulation, such as IgG, IgA or IgM that can bind to the specific bacterial proteins such as the protein A of *S. aureus*, or the protein G of *Streptococcus* with high affinity so that the invading bacteria will be cleared rapidly by the complement/phagocytes from the circulation ^[21–23]^. Our data also showed that there are significantly elevated anti-microbial antibodies in the patients with chronic diseases such as Crohn’s disease (CD) and Sjogren’s syndrome (Sjo) ^[9]^. These chronic “autoimmune” diseases are genetically heterogeneous and polygenic, and the pathogenesis of these diseases is more complex and associated with different types of microbes as demonstrated by the new microbiome studies. The presence of the same anti-microbial antibodies in different clinical diseases such as Crohn’s disease (CD), Sjogren’s syndrome (Sjo) and other autoimmune diseases raises the questions of disease specificity of these anti-microbial antibodies. It remains a possibility that the disease specificity is conferred through genetic susceptibility and genetic heterogeneity to the colonizing microbes within the human body, but not the colonizing microbes themselves, as the commensal microbes are present universally in the human population. Understanding the “point of entry” or the barrier systems including the mucosa and skin as parts of the innate immunity system interacting with the colonizing microbes remains the key ^[24]^. It is also a possibility that the colonizing microbes are undergoing significant changes of virulence factors, becoming pathogenic under unknown circumstances ^[25]^.

Identification of anti-microbial antibodies in patients with chronic diseases but not in normal healthy controls raises questions of bacterial vaccination using the whole bacterial extracts. Historic effort of development of *Mycobacterium tuberculosis* vaccine, like many other bacterial vaccines, was using the whole bacterial extracts or whole bacteria, inactivated or attenuated, and the whole bacterial extract elicits the immune responses and the antibody production in a manner similar to those identified in our patient population, and these antibodies are shown to be highly variable in efficacy to the patients ^[26 27]^. Bacterial components or specific bacterial proteins, however, can be used for vaccination to produce protective, rather than pathogenic antibodies. It seems again that the human body responds to the invading bacteria in specific manners but this specificity of the immune response does not appear conferred through the invading bacteria but the host genetic background, ie, genetic susceptibility determines the host response to the environmental microbes. It should be noted that the viral immunity and viral vaccination is entirely different from those of the bacterial immunity.

Our panel of anti-microbial antibodies appears to be highly sensitive to chronic diseases such as Crohn’s disease (CD) and Sjogren’s syndrome (Sjo). The presence of antimicrobial antibodies in the circulation can separate those with abnormal immune response to the environmental (commensal or pathogenic) microbes from the normal individuals. The value of our panel of anti-microbial antibodies as a diagnostic test to a specific disease type appears to be limited due to the fact that many clinical disease types are found to have elevated anti-microbial antibodies against the same types of microbes in our data. However, the panel is highly sensitive and reliable to identify the individuals with abnormal immune responses to the commensal/pathogenic microbes. These individuals with abnormal immune response to the colonizing microbes can manifest as a variety of disease processes in different organ systems, dependent upon the genetic susceptibility and the phenotypic response of the host. In this regard, the panel test is useful separating those with genetic susceptibility to colonizing microbes from those under environmental stresses or short term infection alone. Furthermore, this panel of anti-microbial antibodies appears to be useful for predicting the occurrence of disease or monitoring the disease progression rather than diagnosis alone, as the abnormal immune response to the microbes can manifest many years earlier than the tissue damage resulting from these immune responses. Longitudinal survey study of normal healthy individuals early with subsequent disease occurrence will answer these interesting questions.

In the new microbiome era it is known that humans are colonized by trillions of diverse microbes and these microbes are commensal and essential in normal human metabolic functions and development ^[10]^. It is also known that there are numerous microbial DNAs within the normal circulation without living microorganisms ^[28]^, and these microbial DNAs, are not eliciting the T-cell/B-cell/plasma cell response for antibody production based on the current literature. How our body recognizes the microbial DNA and proteins within our circulation in relation to health and disease remains to be a research topic for many years.

## References

1. Yu JE, De Ravin SS, Uzel G, et al. High levels of Crohn’s disease-associated anti-microbial antibodies are present and independent of colitis in chronic granulomatous disease. Clin Immunol 2011;138(1):14–22

2. Zholudev A, Zurakowski D, Young W, Leichtner A, Bousvaros A. Serologic testing with ANCA, ASCA, and anti-OmpC in children and young adults with Crohn’s disease and ulcerative colitis: diagnostic value and correlation with disease phenotype. Am J Gastroenterol 2004;99(11):2235–41

3. Landers CJ, Cohavy O, Misra R, et al. Selected loss of tolerance evidenced by Crohn’s disease-associated immune responses to auto-and microbial antigens. Gastroenterology 2002;123(3):689–99

4. Dubinsky MC, Lin YC, Dutridge D, et al. Serum immune responses predict rapid disease progression among children with Crohn’s disease: immune responses predict disease progression. Am J Gastroenterol 2006;101(2):360–7

5. Papp M, Altorjay I, Dotan N, et al. New serological markers for inflammatory bowel disease are associated with earlier age at onset, complicated disease behavior, risk for surgery, and NOD2/CARD15 genotype in a Hungarian IBD cohort. Am J Gastroenterol 2008;103(3):665–81

6. Prideaux L, De Cruz P, Ng SC, Kamm MA. Serological antibodies in inflammatory bowel disease: a systematic review. Inflamm Bowel Dis 2012;18(7):1340–55

7. Zhang P. Anti-microbial antibodies, autoantibodies and autoimmune diseases: Lambert Publishing Company, 2018.

8. Zhang P, Minardi LM, Kuenstner JT, Zekan SM, Kruzelock R. Anti-microbial Antibodies, Host Immunity, and Autoimmune Disease. Front Med (Lausanne) 2018;5:153

9. Zhang P, Minardi, LM., Kuenstner, JT., Zekan, SM., Zhu, F., Hu, YL., Kruzelock, R. Cross–reactivity of antibodies against microbial proteins to human tissues as basis of Crohn’s disease and Sjogren’s syndrome. http://biorxiv.org/content/early/2017/03/13/116574 2017

10. Gilbert JA, Blaser MJ, Caporaso JG, Jansson JK, Lynch SV, Knight R. Current understanding of the human microbiome. Nat Med 2018;24(4):392–400

11. Pascal V, Pozuelo M, Borruel N, et al. A microbial signature for Crohn’s disease. Gut 2017;66(5):813–22

12. Chu H, Khosravi A, Kusumawardhani IP, et al. Gene-microbiota interactions contribute to the pathogenesis of inflammatory bowel disease. Science 2016;352(6289):1116–20

13. Young VB. The role of the microbiome in human health and disease: an introduction for clinicians. Bmj 2017;356:j831

14. Levy M, Thaiss CA, Zeevi D, et al. Microbiota-Modulated Metabolites Shape the Intestinal Microenvironment by Regulating NLRP6 Inflammasome Signaling. Cell 2015;163(6):1428–43

15. Elinav E, Strowig T, Kau AL, et al. NLRP6 inflammasome regulates colonic microbial ecology and risk for colitis. Cell 2011;145(5):745–57

16. Wu HJ, Ivanov, II, Darce J, et al. Gut-residing segmented filamentous bacteria drive autoimmune arthritis via T helper 17 cells. Immunity 2010;32(6):815–27

17. Fridey JL. A compendium of transfusion practice guideline, 2017.

18. Zou KH, O’Malley AJ, Mauri L. Receiver-operating characteristic analysis for evaluating diagnostic tests and predictive models. Circulation 2007;115(5):654–7

19. Olopoenia LA, King AL. Widal agglutination test - 100 years later: still plagued by controversy. Postgrad Med J 2000;76(892):80–4

20. Israel L, Wang Y, Bulek K, et al. Human Adaptive Immunity Rescues an Inborn Error of Innate Immunity. Cell 2017;168(5):789–800 e10

21. Akerstrom B, Bjorck L. Protein L: an immunoglobulin light chain-binding bacterial protein. Characterization of binding and physicochemical properties. J Biol Chem 1989;264(33):19740–6

22. Akerstrom B, Brodin T, Reis K, Bjorck L. Protein G: a powerful tool for binding and detection of monoclonal and polyclonal antibodies. J Immunol 1985;135(4):2589–92

23. Hjelm H, Hjelm K, Sjoquist J. Protein A from Staphylococcus aureus. Its isolation by affinity chromatography and its use as an immunosorbent for isolation of immunoglobulins. FEBS Lett 1972;28(1):73–6

24. Becattini S, Taur Y, Pamer EG. Antibiotic-Induced Changes in the Intestinal Microbiota and Disease. Trends Mol Med 2016;22(6):458–78

25. Granato ET, Harrison F, Kummerli R, Ross-Gillespie A. Do Bacterial “Virulence Factors” Always Increase Virulence? A Meta-Analysis of Pyoverdine Production in Pseudomonas aeruginosa As a Test Case. Front Microbiol 2016;7:1952

26. Aronson NE, Santosham M, Comstock GW, et al. Long-term efficacy of BCG vaccine in American Indians and Alaska Natives: A 60-year follow-up study. Jama 2004;291(17):2086–91

27. Roy A, Eisenhut M, Harris RJ, et al. Effect of BCG vaccination against Mycobacterium tuberculosis infection in children: systematic review and metaanalysis. Bmj 2014;349:g4643

28. Kowarsky M, Camunas-Soler J, Kertesz M, et al. Numerous uncharacterized and highly divergent microbes which colonize humans are revealed by circulating cell-free DNA. Proc Natl Acad Sci U S A 2017;114(36):9623–28

